# Accuracy of whole-genome sequence imputation using hybrid peeling in large pedigreed livestock populations

**DOI:** 10.1101/771576

**Authors:** Roger Ros-Freixedes, Andrew Whalen, Ching-Yi Chen, Gregor Gorjanc, William O Herring, Alan J Mileham, John M Hickey

**Author notes:** Corresponding author: RRF. Email addresses: AW, CYC, GG, WOH, AJM, JMH.

## Abstract

**Background:** We demonstrate high accuracy of whole-genome sequence imputation in large livestock populations where only a small fraction of individuals (2%) had been sequenced, mostly at low coverage.

**Methods:** We used data from four pig populations of different sizes (18,349 to 107,815 individuals) that were broadly genotyped at densities between 15,000 and 75,000 markers genome-wide. Around 2% of the individuals in each population were sequenced (most at 1x or 2x and a small fraction at 30x; average coverage per individual: 4x). We imputed whole-genome sequence with hybrid peeling. We evaluated the imputation accuracy by removing the sequence data of a total of 284 individuals that had been sequenced at high coverage, using a leave-one-out design. We complemented these results with simulated data that mimicked the sequencing strategy used in the real populations to quantify the factors that affected the individual-wise and variant-wise imputation accuracies using regression trees.

**Results:** Imputation accuracy was high for the majority of individuals in all four populations (median individual-wise correlation was 0.97). Individuals in the earliest generations of each population had lower accuracy than the rest, likely due to the lack of marker array data for themselves and their ancestors. The main factors that determined the individual-wise imputation accuracy were the genotyping status of the individual, the availability of marker array data for immediate ancestors, and the degree of connectedness of an individual to the rest of the population, but sequencing coverage had no effect. The main factors that determined variant-wise imputation accuracy were the minor allele frequency and the number of individuals with sequencing coverage at each variant site. These results were validated with the empirical observations.

**Conclusions:** The coupling of an appropriate sequencing strategy and imputation method, such as described and validated here, is a powerful strategy for generating whole-genome sequence data in large pedigreed populations with high accuracy. This is a critical step for the successful implementation of whole-genome sequence data for genomic predictions and fine-mapping of causal variants.

## Background

In this paper we demonstrate high accuracy of whole-genome sequence imputation in large livestock populations where only a small fraction of individuals (2%) had been sequenced, mostly at low coverage. Using data from pig populations we show that imputation accuracy was very high for individuals that were genotyped with marker arrays with densities that ranged between 15,000 and 75,000 markers genome-wide. We also used simulations to quantify the factors that determined the imputation accuracy achieved for each individual and variant, validated those results with the empirical observations from real data, and performed robustness tests to determine the impact of data misassignment and pedigree errors on the imputation accuracy.

Sequence data has the potential to empower the identification of causal variants that underlie quantitative traits or diseases [1–4], enhance livestock breeding [5–7], and increase the precision and scope of population genetic studies [8,9]. For sequence data to be used routinely in research and breeding, low-cost sequencing strategies must be deployed in order to assemble large data sets that capture most of the sequence diversity in a population and enable harnessing of its potential. One possible strategy is to sequence a subset of the individuals in a population at low coverage and then to perform imputation of whole-genome sequence data for the remaining individuals [10–12].

Such a strategy is likely to perform well in livestock breeding populations, where individuals have a high degree of relatedness, allowing low-coverage sequence data to be pooled across individuals that share a haplotype and imputed to individuals who share that haplotype and have small amounts of sequenced data or who do not have any sequence data. Due to the implementation of genomic selection in livestock breeding populations, many individuals in breeding nucleus populations have already been genotyped with marker arrays. This genotype data can be used to identify the individuals that share haplotype segments and to select individuals for sequencing that will be more informative from an imputation perspective given a limited budget [13,14].

We have recently proposed ‘hybrid peeling’ [15], a fast and accurate imputation method explicitly designed for jointly calling, phasing and imputing whole-genome sequence data in large and complex multi-generational pedigreed populations where individuals can be sequenced at variable coverage or not sequenced at all. Hybrid peeling is a two-step process. In the first step, multi-locus iterative peeling is performed to estimate the segregation probabilities for a subset of segregating sites (e.g., the markers on a genotyping array). In the second step, the segregation probabilities are used to perform fast single-locus iterative peeling on every segregating site discovered in the genome. This two-step process allows the computationally demanding multi-locus peeling step to be performed on only a subset of the variants, while still leveraging linkage information for the remaining variants.

These properties make hybrid peeling a very appealing imputation method for the cost-effective generation of whole-genome sequence data for large pedigreed populations that have already been extensively genotyped using marker arrays and in which a small proportion of the individuals have been sequenced with variable coverage. In the situations described, the sequence data will be sparsely distributed across the pedigree and there may be great variability in the amount of data to which each individual is exposed. Understanding which factors affect individual-wise and variant-wise imputation accuracy and how their effects are mediated is important for determining how this sequencing strategy, together with hybrid peeling, performs in real settings that are common in animal breeding and for enabling accuracy-aware quality control of the imputed data before downstream analyses. Such knowledge could be used in the future to design cost-effective routine whole-genome sequencing strategies.

The objectives of this study were to: (i) demonstrate if whole-genome sequence data could be imputed with high accuracy in a variety of pig pedigrees when small subsets of individuals are sequenced, mostly at low coverage; (ii) quantify the factors that determine the individual-wise and variant-wise imputation accuracy; and (iii) quantify the impact of data misassignment and pedigree errors on imputation accuracy. Our results showed that high overall imputation accuracies can be achieved for whole-genome sequence data in large pedigreed populations using hybrid peeling provided that the individuals are connected to a sufficient number of informative relatives with marker array or sequence data.

## Materials and Methods

We structured the study in three tests. In Test 1 we evaluated the imputation accuracy of hybrid peeling in four populations of different sizes by removing the sequence data of 284 individuals that had been sequenced at high coverage, using a leave-one-out validation design. In Test 2 we used simulated data based on three other real pedigrees to quantify which factors determined the individual-wise and variant-wise imputation accuracy of hybrid peeling with regression trees. We used simulated data to provide a much larger sample size where the true genotypes were known, and we then used the observations in the real data to validate the findings. In Test 3, we evaluated the potential impact that data misassignment and pedigree errors could potentially have on the imputation accuracy by introducing deliberate errors in the real data. In what follows we first describe how the data was generated and then how the different tests were performed.

### Real data

#### Populations and sequencing strategy

We performed whole-genome sequencing of 4,427 individuals from four commercial pig breeding lines (Genus PIC, Hendersonville, TN) using a total coverage of approximately 18,514x. The populations selected for this study differed in size, and approximately 2% (1.7-2.5%) of the individuals in each population were sequenced, mostly at low coverage. The first population had 18,349 (20k) individuals and 445 of these were sequenced with a total coverage of 1,852x. The second population had 34,425 (35k) individuals and 760 of these were sequenced with a total coverage of 3,192x. The third population had 68,777 (70k) individuals and 1,366 of these were sequenced with a total coverage of 5,280x. The fourth population had 107,815 (110k) individuals and 1,856 of these were sequenced with a total coverage of 8,190x. We sorted the pedigrees of each population so that parents appeared before their progeny. Thus, relative position in the pedigree was used as a proxy for the generation to which an individual belonged.

We selected the individuals and the coverage at which they were sequenced using a three-step strategy: (1) we first selected sires and dams that contributed most genotyped progeny in the pedigree (referred to as ‘top sires and dams’) to be respectively sequenced at 2x and 1x; (2) conditional on the first step, we used AlphaSeqOpt part 1 [13] to identify the individuals whose haplotypes represented the greatest proportion of the population haplotypes (referred to as ‘focal individuals’) and to determine an optimal level of sequencing coverage between 0x and 30x for these individuals and their immediate ancestors (i.e., parents and grandparents) under a total cost constraint; and (3) conditional on the second step, we used the AlphaSeqOpt part 2 [14] to identify individuals that carried haplotypes whose cumulative coverage was low (i.e., below 10x) and distributed 1x sequencing amongst those individuals so that the cumulative coverage on the haplotypes could be increased (i.e., at or above 10x). AlphaSeqOpt used haplotypes inferred from marker array genotypes (GGP-Porcine HD BeadChip; GeneSeek, Lincoln, NE), which were phased with AlphaPhase [16] and imputed with AlphaImpute [17]. The sequencing resources were split so that approximately 30% of the sequencing resources were used for sequencing the top sires at 2x, 15% for the top dams at 1x, 25% for the focal individuals and their immediate ancestors at variable coverage [13], and the remaining 30% for individuals that carried under-sequenced haplotypes at 1x [14]. In step 2 we identified a total of 284 individuals across the four populations who were sequenced at high coverage (15x or 30x). Of these, 37 belonged to the 20k population, 65 to the 35k population, 92 to the 70k population, and 90 to the 110k population. Many of these individuals belonged to early generations of the pedigree of each population. The rest of the sequenced individuals were sequenced at low coverage (1x, 2x or 5x). The number of individuals sequenced and the coverage at which they were sequenced is summarized for each population in Table 1.

**Table 1.** Distribution of sequencing coverages by population.

| Population | Individuals<br>sequenced | Individuals sequenced by coverage |  |  |  | Total<br>coverage |
| --- | --- | --- | --- | --- | --- | --- |
|  |  | 1x | 2x | 5x | 15-30x |  |
| 20k | 445 | 217 | 176 | 15 | 37 | 1,852x |
| 35k | 760 | 394 | 274 | 27 | 65 | 3,192x |
| 70k | 1,366 | 685 | 545 | 44 | 92 | 5,280x |
| 110k | 1,856 | 1,044 | 649 | 73 | 90 | 8,190x |

#### Sequencing and data processing

Tissue samples were collected from ear punches or tail clippings. Genomic DNA was extracted using Qiagen DNeasy 96 Blood & Tissue kits (Qiagen Ltd., Mississauga, ON, Canada). Paired-end library preparation was conducted using the TruSeq DNA PCR-free protocol (Illumina, San Diego, CA). Libraries for sequencing at low coverage (1x to 5x) were produced with an average insert size of 350 base pairs and sequenced on a HiSeq 4000 instrument (Illumina, San Diego, CA). Libraries for sequencing at high coverage (15x or 30x) were produced with an average insert size of 550 base pairs and sequenced on a HiSeq X instrument (Illumina, San Diego, CA). All libraries were sequenced at Edinburgh Genomics (Edinburgh Genomics, University of Edinburgh, Edinburgh, UK). Most pigs were also genotyped either at low density (LD; 15,000 markers) using the GGP-Porcine LD BeadChip (GeneSeek, Lincoln, NE) or at high density (HD; 75,000 markers) using the GGP-Porcine HD BeadChip (GeneSeek, Lincoln, NE).

DNA sequence reads were pre-processed using Trimmomatic [18] to remove adapter sequences from the reads. The reads were then aligned to the reference genome *Sscrofa11.1* (GenBank accession: GCA_000003025.6; [19]) using the BWA-MEM algorithm [20]. Duplicates were marked with Picard (http://broadinstitute.github.io/picard). Single nucleotide polymorphisms (**SNPs**) and short insertions and deletions (indels) were identified with the variant caller GATK HaplotypeCaller (GATK 3.8.0; [21,22]) using default settings. Variant discovery with GATK HaplotypeCaller was performed separately for each individual. A joint variant set for all the individuals in each population was obtained by extracting the variant sites from all the individuals. Between 20 and 30 million variants were discovered in each population.

To avoid biases towards the reference allele introduced by GATK when applied on low-coverage sequence data we extracted the read counts supporting each allele directly from the aligned reads stored in the BAM files with a pile-up function using the pipeline described in [23]. This pipeline uses the tool pysam (version 0.13.0; https://github.com/pysam-developers/pysam), which is a wrapper around htslib and the samtools package [24]. We extracted the read counts for all biallelic SNP positions, after filtering out variants with mean coverage 3 times greater than the average realized coverage (considered as indicative of potential repetitive regions) with VCFtools [25].

We performed additional quality control on the pedigree by determining the number of Mendelian inconsistencies (percentage of opposing homozygous) between each parent-progeny pair. We applied the following criteria: (1) we removed marker array or sequence data of an individual, when the genotype data was incompatible with that of all its available parents and progeny (this was done because it could indicate data misassignment for that individual); (2) we removed parent-progeny pedigree links when the genotype data available was incompatible for only a pair of individuals but not for their other parents and progeny; and (3) we created a dummy parent with no genotype data when the genotype data of a group of littermates was incompatible with one of its parents but both the parent and the littermates were not incompatible with the rest of their parents and progeny (this was done to preserve the full-sib relationship between those individuals).

### Simulated data

In order to test the factors that influenced imputation accuracy, we simulated genetic data for three populations of different sizes: 15,187 (15k), 29,974 (30k), and 64,598 (65k) individuals. The pedigrees of these populations were a subset of the real pedigrees of the 20k, 35k, and 110k populations used for the analyses of real data. As in the analyses of real data, the pedigrees were sorted so that parents appeared before their progeny. Genomic data for each population was simulated using the software AlphaSim [26]. Each simulation was repeated twice and results were averaged across repetitions. Below, we present only a brief description of the simulation strategy. The full details of the simulation are described in a companion paper [27].

Genomic data were simulated for 20 chromosomes, each 100 cM in length. A total of 150,000 SNPs per chromosome (3 million SNPs genome-wide) were simulated in order to represent whole-genome sequence. A subset of 3,000 SNPs per chromosome (60,000 SNPs genome-wide) was used as a high-density marker array (HD). A smaller subset of 300 SNPs per chromosome (6,000 SNPs genome-wide) nested within the high-density marker array was used as a low-density marker array (LD). Each individual was assigned HD or LD marker array data based on the density at which they were genotyped in real data. The sequence read counts for each individual and SNP were simulated by sampling sequence reads using a Poisson-gamma model that gave variable sequenceability at each SNP and variable number of reads for each individual at each SNP [28,29].

The individuals to be sequenced and their sequencing coverage were selected using a combination of pedigree-and haplotype-based methods that emulated the sequencing strategy that was used for the real data. In implementing this approach, for simplicity the simulated sequencing resources were split in an equitable way so that 25% of the sequencing resources were used for sequencing top sires at 2x, 25% for top dams at 1x, 25% for the focal individuals and their immediate ancestors at variable coverage [13], and the remaining 25% for individuals that carried under-sequenced haplotypes at 1x [14]. The total level of investment for sequencing was equivalent to the cost of sequencing 2% of the population at 2x, and thus resulted in a similar number of sequenced individuals as in the real data.

### Imputation using hybrid peeling

Imputation was performed in each population separately using hybrid peeling, as implemented in AlphaPeel [15] with the default settings. Hybrid peeling extends the methods of Kerr and Kinghorn [30] for single-locus iterative peeling and of Meuwissen and Goddard [31] for multi-locus iterative peeling to efficiently call, phase and impute whole-genome sequence data in complex multi-generational pedigrees with loops. Multi-locus iterative peeling was performed on all available marker array data to estimate the segregation probabilities for each individual. The individuals genotyped with LD marker arrays were not imputed to HD prior to this step. The segregation probabilities were used for segregation-aware single-locus iterative peeling for the remaining segregating variants.

### Imputation accuracy tests

#### Test 1: Imputation accuracy in populations of different size

The imputation accuracy in the real data was estimated using a leave-one-out design. In each leave-one-out round, hybrid peeling was performed after removing the sequence data of one of the 284 individuals that were sequenced at high coverage (either 15 or 30x) in the four populations, which produced a total of 284 validation rounds across the four populations. We used the genotypes imputed for these individuals using the full data as the true genotypes. To reduce computational requirements, accuracy was only assessed on a subset of 50,000 non-consecutive SNPs on a single chromosome. The chromosome that we used was chromosome 5, which was selected randomly and has an intermediate size compared to the other pig chromosomes. Tests in other chromosomes gave similar results. The 50,000 variants that we tested included all the markers from the arrays that map to this chromosome (∼3,000), while the rest were chosen randomly from sequence variants discovered along the chromosome.

We measured individual-wise and variant-wise imputation accuracy with the genotype concordance, measured as the percentage of correct genotypes, and the correlation between the true genotypes and imputed dosages. The individual-wise correlation was calculated after correcting for minor allele frequency (MAF), as recommended by Calus et al. [32]. In the context of this study, we found that the relationship between the raw correlation uncorrected for MAF and the dosage corrected for MAF was nearly linear (see Figure S1). To facilitate comparison with other studies that report the uncorrected (raw) allele dosage correlations, we found that the MAF corrected correlations of 0.75, 0.80, 0.85, 0.90, and 0.95 were respectively equivalent to the raw correlations of 0.89, 0.91, 0.93, 0.96, and 0.98. The variant-wise imputation accuracy was measured as the correlation between the imputed allele dosages and true genotypes without any correction.

#### Test 2: Factors that affect individual-wise and variant-wise imputation accuracy

In this test we assessed the factors that influenced imputation accuracy in the simulated data. The simulated data was used to provide a much larger sample size where the true genotypes were known. Just as for the real data, we ran single-locus peeling only on a random subset of SNPs among all sequence variants; in this case on a total of 5,000 non-consecutive SNPs taken from across three chromosomes to reduce computational requirements, although the full set of 20 chromosomes were simulated to represent realistic genetic architecture and haplotype diversity, which was needed to ensure that the properties of AlphaSeqOpt, which is a haplotype-based method, matched those of the of real data. We assessed the factors that influenced imputation accuracy by building regression trees. The regression trees were built using the data from 219,518 simulated individuals and a total of 30,000 variants (5,000 variants from each population and replicate).

The regression tree for the individual-wise imputation accuracy was based on the amount of information that was available for the individual itself and its close relatives (4 relationship levels: grandparents, parents, progeny, and grandprogeny). The factors included: (i) size of the population to which they belonged (15k, 30k, or 65k individuals); (ii) marker array density of the individual (3 genotyping statuses: not genotyped, genotyped at LD, or genotyped at HD); (iii) number of close relatives that were genotyped at each genotyping density (12 variables; 4 relationship levels and 3 genotyping statuses); (iv) sequencing coverage of the individual; (v) number of close relatives that were sequenced and their cumulative sequencing coverage (8 variables; 2 variables for each of the 4 relationship levels); and (vi) connectedness to the population, which was measured as the sum of coefficients of relationship between an individual and the rest of individuals in the pedigree. The regression tree was built using the ‘rpart’ R package [33], allowing partitions that increased the overall R^2^ by 0.005 at each step. Consecutive binary partitions based on the same variable were considered as multi-part.

The factors in the regression tree for the variant-wise imputation accuracy included: (i) size of the population (15k, 30k, or 65k individuals); (ii) MAF; (iii) relative position of the variant within a chromosome; (iv) distance of a variant to the nearest variant from the marker array (this distance was 0 if that variant was present on the marker array); (v) cumulative sequencing coverage across individuals at that variant site; and (vi) number of individuals with at least one sequencing read covering that variant site. As with the individual-wise imputation accuracy, we allowed partitions that increased the overall R^2^ by 0.005 at each step and consecutive binary partitions based on the same variable were considered as multi-part.

We then used the 284 high-coverage individuals in the real data for validation, by comparing the results of the regression trees from the simulated data with the imputation accuracies observed in the real data. A regression tree was not separately created for the real data due to the small number of high-coverage individuals in our validation set. To further assess which factors affected the individual-wise imputation accuracy in the real data we fitted a linear model predicting imputation accuracy against each of the factors used for the regression tree.

#### Test 3: Impact of data misassignment and pedigree errors

We tested the impact that data misassignment and pedigree errors could have on the imputation results by introducing deliberate errors to the real data. We considered three types of errors: sequence data misassignment, marker array data misassignment, and pedigree errors. For each type of error we created 284 scenarios, in which we altered the data of each of the individuals that were sequenced at high coverage in each population, one at a time. The three types of errors were defined as follows, to represent some worst-case scenarios:

- *Sequence data misassignment.* We replaced the sequence data of the target individual by that of a random individual from the same population that had been sequenced at high coverage.

- *Marker array data misassignment.* We replaced the marker array data of the target individual by that of a random individual from the same population that had been genotyped at HD, regardless of its own genotyping status or density.

- *Pedigree errors.* We assigned a random progeny from one of the individuals sequenced at high coverage from the same population to the target individual.

The impact of the data misassignment and pedigree errors on imputation accuracy was measured as the correlation between the allele dosages using the correct data and the erroneous data. The impact of these errors was assessed on the target individual where the error was introduced but also on its grandparents, parents, progeny, and grandprogeny to evaluate how the errors could propagate to relatives of the target individual. In the case of the pedigree errors we also assessed the impact of the pedigree error on the misassigned progeny and grandprogeny. As a control we also assessed the allele dosage correlation on the target individual and its relatives when the data of the target individual was removed, as performed in Test 1.

## Results

### Imputation accuracy in populations of different size

#### Individual-wise imputation accuracy

The imputation accuracy in the real data was high for most of the tested individuals. The average individual-wise dosage correlation was 0.94 but there was substantial variation with an asymmetrical distribution (median: 0.97; min: 0.11; max: 1; interquartile range: 0.94-0.98). The average individual-wise genotype concordance was 97.1% (median: 98.4%; min: 78.9%; max: 100%; interquartile range: 97.1-98.9%). Some of the oldest individuals that belonged to the earliest generations of the pedigree (some of the 106 individuals located in the first 20% of the pedigree) had lower imputation accuracy than individuals in the remainder of pedigree, who had consistently high imputation accuracy. This pattern was observed for all four populations. Figure 1 shows the imputation accuracy, measured as the individual-wise dosage correlation, plotted against relative position in the pedigree, the marker array density of the individual, or size of the population to which they belonged. Figure 2 shows the same but with imputation accuracy measured as the individual-wise genotype concordance. The imputation accuracy of the individuals in later generations (the 178 individuals after the first 20% of the pedigree) was higher (Figures S2 and S3), with an average dosage correlation of 0.97 and with much lower variability (median: 0.98; min: 0.69; max: 1; interquartile range: 0.96-0.99), and an average genotype concordance of 98.3% (median: 98.7%; min: 86.9%; max: 100%; interquartile range: 98.3-99.0%).

**Figure 1.**
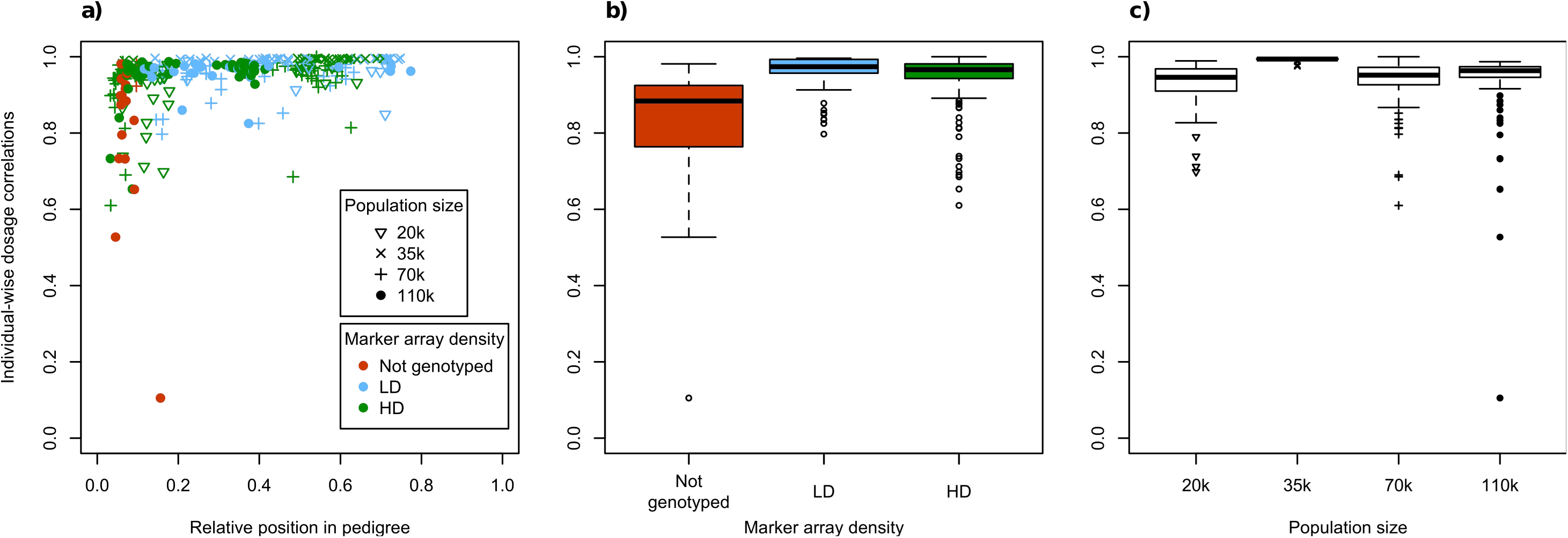
Individual-wise dosage correlation in the real data with respect to (a) relative position of the tested individuals within a pedigree, (b) genotyping marker array density, and (c) population size.

**Figure 2.**
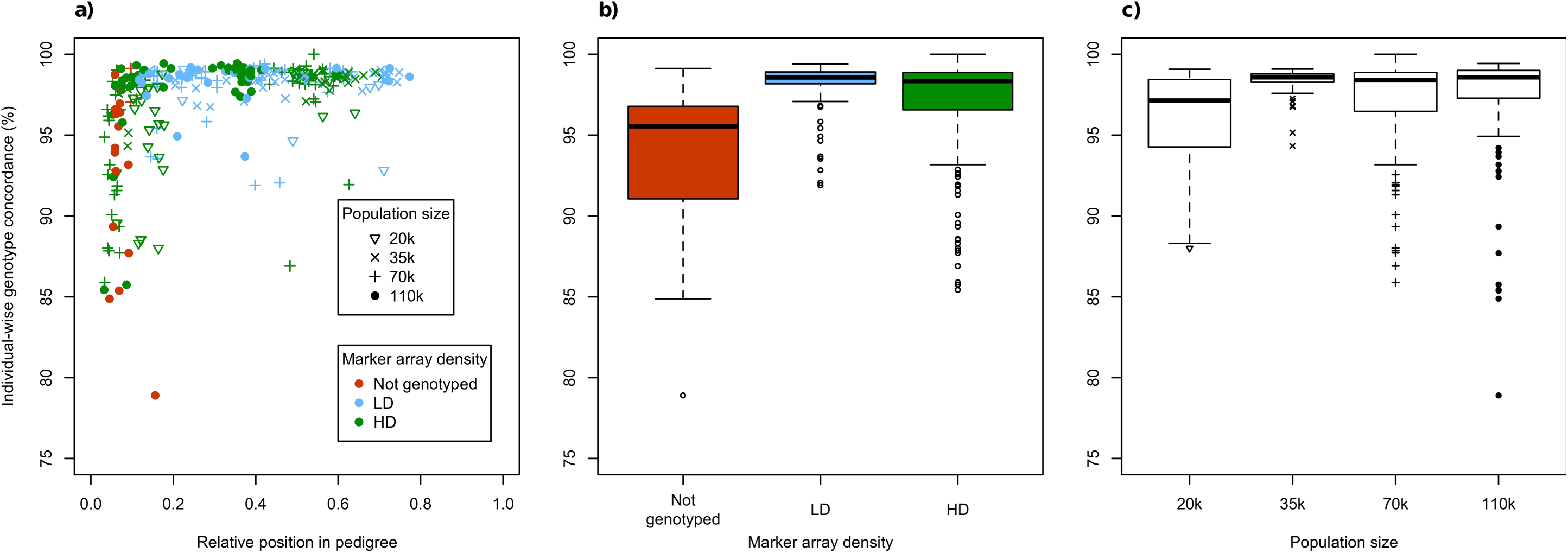
Individual-wise genotype concordance in the real data with respect to (a) relative position of the tested individuals within a pedigree, (b) genotyping marker array density, and (c) population size.

The marker array density of the individuals was confounded with the number of ancestors that were genotyped with marker arrays. The non-genotyped individuals (n=19) and approximately half of the individuals genotyped at HD (n=87 out of 157) belonged to early generations of the pedigree (Figures 1a and 2a), which reduced the chances that they had ancestors with data and penalized the imputation accuracy for these two groups of individuals (Figures 1b and 2b). On the contrary, most individuals genotyped at LD belonged to later generations (n=91 out of 108), ensuring that their ancestors had enough data to enable high imputation accuracies for the LD individuals. The average dosage correlation for the non-genotyped individuals was 0.81, for the HD individuals was 0.94, and for the LD individuals was 0.96. The average dosage correlation for the HD individuals in the earliest generations was lower (0.91) than for the HD individuals in later generations (0.97). For individuals in the later generations there were no significant differences between marker array densities and the average dosage correlation of both the HD and LD individuals was 0.97 (Figures S2b and S3b). There was no clear trend that population size affected imputation accuracy (Figures 1c and 2c), especially for individuals in the later generations (Figures S2c and S3c). The population with 35k individuals had higher imputation accuracy than the other three populations but this was more likely due to population-specific characteristics, related to unbalanced distributions of the tested individuals across generations and genotyping statuses or potentially to pedigree structure, rather than population size. The 35k population had only 5 out of 65 individuals in the first 20% of the pedigree, compared to a much greater proportion in the other populations (from 15 out of 37 in the 15k population to 56 out of 92 in the 65k population).

#### Variant-wise imputation accuracy

The variant-wise imputation accuracy was also high. The average variant-wise dosage correlation was 0.88 (median: 0.96; min: −0.33; max: 1; interquartile range: 0.92-0.99) and the average variant-wise genotype concordance was 96.3% (median: 97.8%; min: 24.3%; max: 100%; interquartile range: 95.4-100%). Variant-wise dosage correlations were much higher when the individuals from the first 20% of the pedigree, which had lower individual-wise imputation accuracy, were excluded from the calculation. The average variant-wise dosage correlation calculated from the 178 individuals after the first 20% of the pedigree was 0.93 (median: >0.99; min: −0.46; max: 1; interquartile range: 0.97-1) and the average variant-wise genotype concordance was 97.4% (median: 100%; min: 13.3%; max: 100%; interquartile range: 97.9-100%).

Variant-wise imputation accuracy was lower for low-frequency variants, compared to more common variants. Figure 3 shows the distribution of the dosage correlation for variants across the MAF spectrum. The only MAF category where the average dosage correlation decreased when the individuals from the first 20% of the pedigree were excluded was for MAF≤0.001 (Figure 3b), likely because the individuals in the early generations were biased towards the major allele, which would inflate imputation accuracy for low MAF variants (the major allele is more likely to be true). Figure S4 shows the distribution of the genotype concordance for variants across the MAF spectrum. Note that the genotype concordance increases at lower MAF because the probability that the true genotype is the most common one increases, highlighting that genotype concordance is misleading as a measure of imputation accuracy and thus should be interpreted with care [34,35].

**Figure 3.**
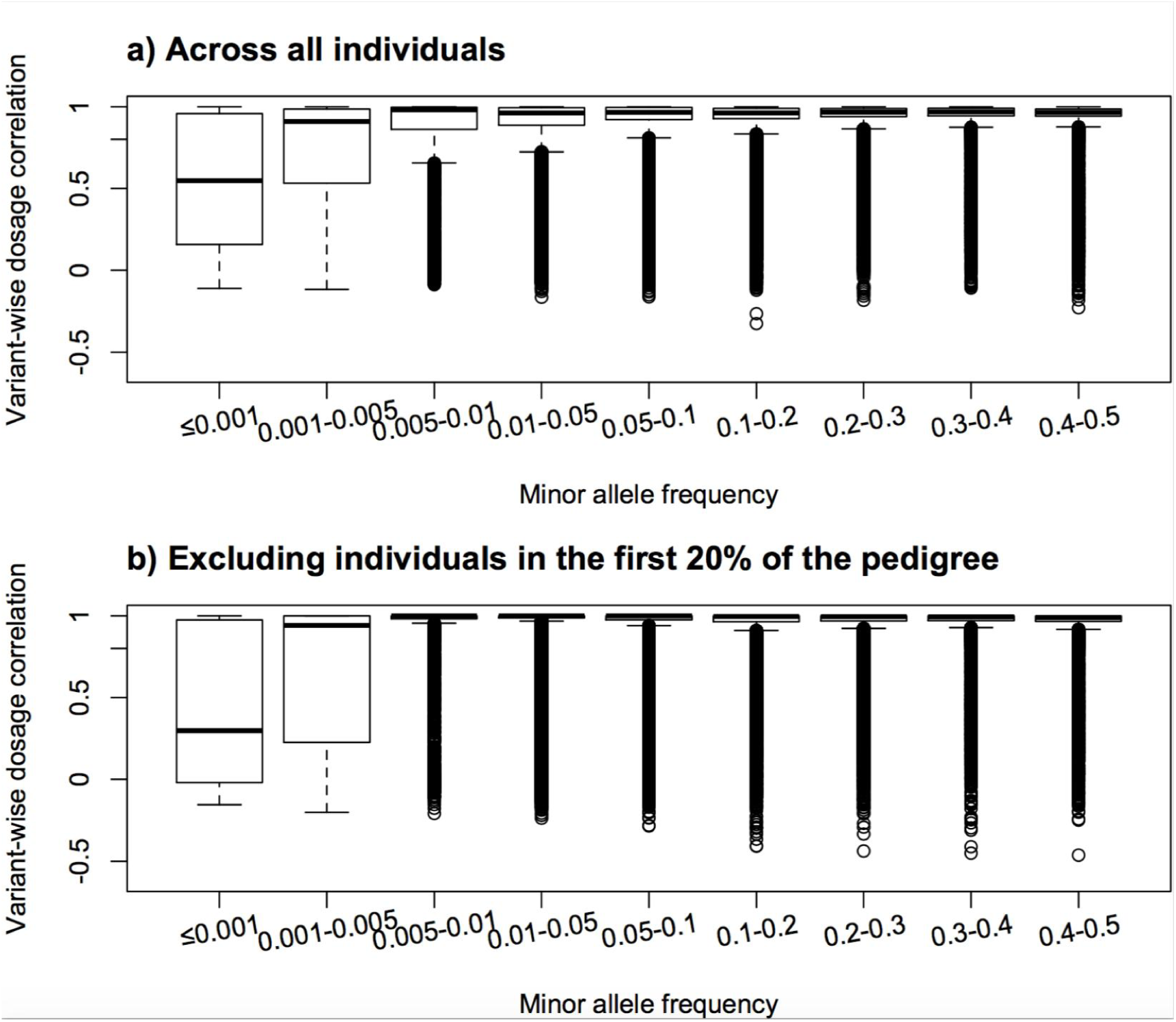
Variant-wise dosage correlation in the real data with respect to minor allele frequency. Results are shown for (a) all individuals or (b) after excluding the individuals in the first 20% of the pedigree.

### Factors that affect individual-wise imputation accuracy

The main factors that determined individual-wise imputation accuracy were whether the individual itself was genotyped with a marker array, the number of close relatives of that individual that were genotyped with a marker array (primarily parents and grandparents), and the connectedness of that individual to the rest of the population. The number of close relatives of an individual that were sequenced was a significant factor for the imputation accuracy of the 284 tested individuals in a linear model, but only the number of sequenced parents or progeny were influential partitioning factors in the regression trees based on the simulated data. The sequencing status of the individual itself or the sequencing coverage of its relatives were not influential partitioning factors in the regression trees. The results were consistent between the simulated and the real data.

The regression tree for the factors that affect individual-wise dosage correlations in the simulated data is shown in Figure 4a. The first partitioning factor was the availability of marker array data of the grandparents. Individuals without genotyped grandparents had much lower imputation accuracy (0.47, n=10,794) than individuals with at least one genotyped grandparent (0.96, n=208,724). In contrast, the number of genotyped parents was not an influential partitioning factor. A likely explanation for this observation was that the number of genotyped grandparents and the number of genotyped parents in the populations were confounded. Specifically, if an individual had genotyped grandparents it was likely that it also had genotyped parents because non-genotyped grandparents were likely to be individuals from very early generations (e.g., the base generation) and most individuals with progeny in subsequent generations were genotyped. For individuals without genotyped grandparents, other sources of information from the ancestors, such as availability of any sequenced parents, increased their imputation accuracy from 0.40 (n=7,516) to 0.63 (n=3,278). After these initial partitions, the next partitioning factor was whether or not the individual itself was genotyped with a marker array, regardless of marker array density. This partition revealed an asymmetry in that individuals without genotyped grandparents or with only one genotyped grandparent were mostly not genotyped themselves (n=6,877 out of 7,516 individuals without any genotype data from their ancestors), whereas the individuals with genotyped grandparents were mostly genotyped (n=194,104 out of 208,724 individuals with genotyped grandparents). For non-genotyped individuals, having some genotyped or sequenced progeny and grandprogeny improved their imputation accuracy. For genotyped individuals, regardless of genotyping density, connectedness to the rest of the population was the main factor that determined imputation accuracy, with the dosage correlation increasing with connectedness from 0.89 (n=9,446) to 0.98 (n=184,658).

**Figure 4.**
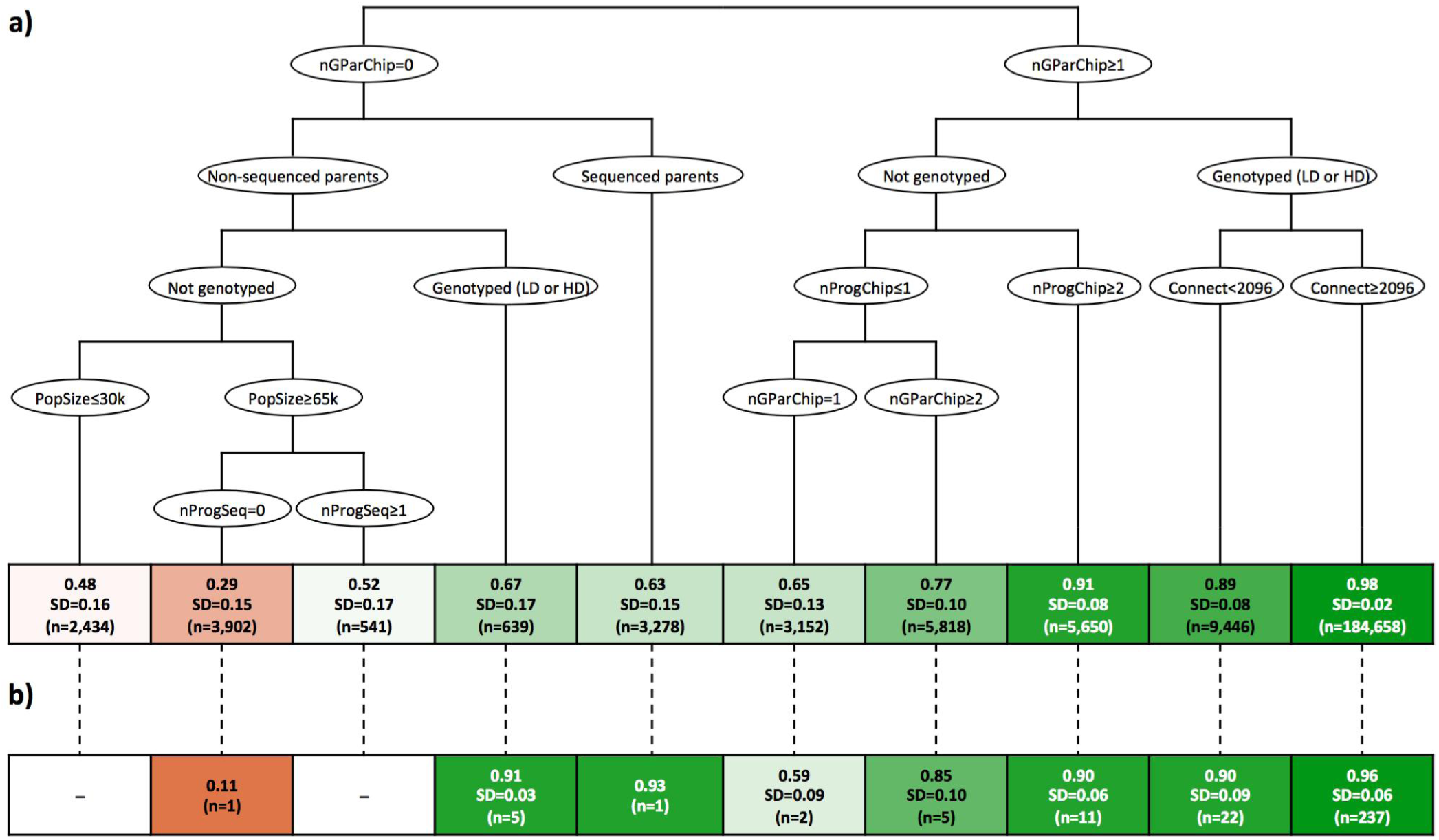
Regression tree of the factors that affected individual-wise dosage correlation in (a) the simulated data and (b) comparison to the real data. Variables include genotype status, number of grandparents genotyped with marker array (nGParChip), number of progeny genotyped with marker array (nProgChip), number of sequenced progeny (nProgSeq), connectedness to the rest of the population (Connect), and population size (PopSize).

The regression tree for the factors that affect individual-wise genotype concordance in the simulated data is shown in Figure 5a. It had a similar pattern to that observed for the dosage correlation. The first partitioning factor was whether or not the individual itself was genotyped with a marker array. For non-genotyped individuals, the next partitioning factors were the availability of marker array data of the grandparents, the parents (if none or only one grandparent were genotyped), and progeny. As the number of genotyped close relatives increased, genotype concordances of the non-genotyped individuals increased from 69.6% (n=8,834) to 93.7% (n=5,022). For genotyped individuals, the next partitioning factor was the connectedness to the rest of the population. The genotype concordance increased with connectedness from 88.4% (n=6,680) to 98.6% (n=152,322). In individuals with low connectedness, availability of marker array data of the grandparents helped improve their genotype concordance.

**Figure 5.**
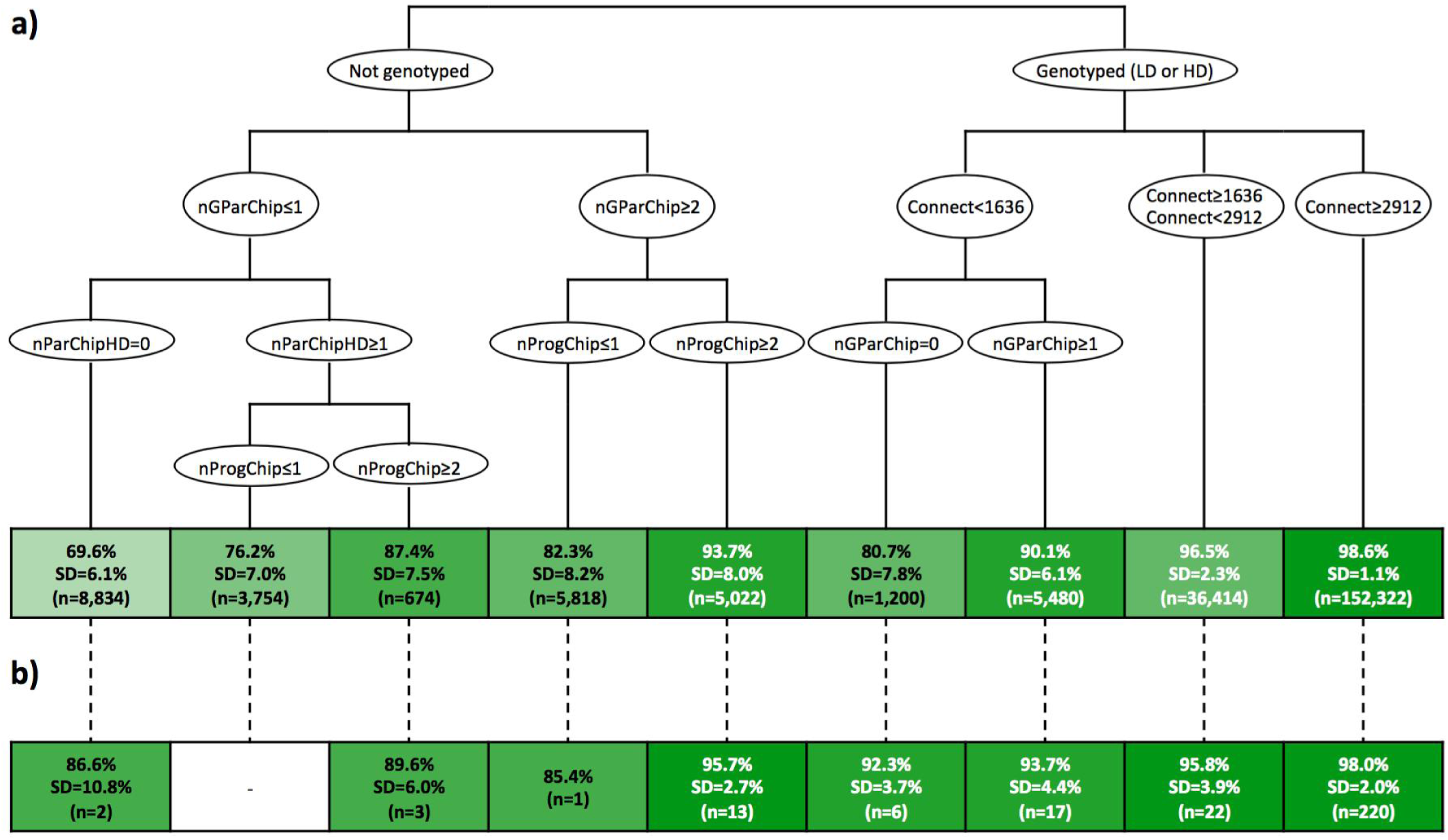
Regression tree of the factors that affected individual-wise genotype concordance in (a) the simulated data and (b) comparison to the real data. Variables include genotype status, number of grandparents genotyped with marker array (nGParChip), number of parents genotyped with high-density marker array (nParChipHD), number of progeny genotyped with marker array (nProgChip), number of grandprogeny genotyped with marker array (nGProgChip), and connectedness to the rest of the population (Connect).

The dosage correlations and genotype concordances observed in the real data were consistent with the partitions of the regression tree based on the simulated data (Figures 4b and 5b). The analysis of the factors that affected the individual-wise imputation accuracy observed in the real data with a linear model largely supported these patterns. Table 2 summarises the factors that were significantly associated with individual-wise imputation accuracy when measured as dosage correlations or genotype concordances. Broadly, the significant factors were the same for both measures of imputation accuracy. The significant factors included the number of genotyped ancestors, but not the number of genotyped descendants, and the number of sequenced relatives, but generally not their cumulative sequencing coverage. The number of parents genotyped with marker arrays at both LD and HD were generally significant factors (*p*-value≤0.001). The number of grandparents genotyped was also significant at HD (*p*-value≤0.016) but not at LD (*p*-value≥0.614). The number of genotyped progeny and grandprogeny were not significant factors (*p*-value≥0.062). The number of sequenced ancestors and descendants were also significant factors (*p*-value≤0.016). The cumulative sequencing coverage of the parents and grandprogeny was significant (*p*-value=0.016 to 0.044) but not that of the grandparents and progeny (*p*-value≥0.100). The factors that referred to the amount of information available for the individuals themselves were also significant, including both their genotyping status (*p*-value≤0.001) and their connectedness to the rest of the population (*p*-value≤0.031). However, the marker array density was confounded with the generation to which the individuals belonged and, therefore, with the number of ancestors that were genotyped with marker arrays (Figure 1). Population size was also a significant factor (*p*-value≤0.001), but likely confounded with population-specific factors (Figure 1).

**Table 2.** Factors that affect individual-wise imputation accuracy on the real data (*p*-value).

| Factor | Allele dosage correlation | Genotype concordance |
| --- | --- | --- |
| Population size | <0.001 *** | <0.001 *** |
| Individual data |  |  |
| Genotyping status | <0.001 *** | <0.001 *** |
| Connectedness to the rest of population | 0.031 * | <0.001 *** |
| Number of relatives genotyped with marker array |  |  |
| Grandparents at LD<sup>a</sup> | 0.707 | 0.614 |
| Grandparents at HD<sup>a</sup> | 0.016 * | <0.001 *** |
| Parents at LD | 0.059 | <0.001 *** |
| Parents at HD | <0.001 *** | <0.001 *** |
| Progeny at LD | 0.062 | 0.202 |
| Progeny at HD | 0.553 | 0.314 |
| Grandprogeny at LD | 0.926 | 0.899 |
| Grandprogeny at HD | 0.996 | 0.681 |
| Number of relatives sequenced |  |  |
| Grandparents | 0.003 ** | <0.001 *** |
| Parents | <0.001 *** | <0.001 *** |
| Progeny | 0.002 ** | <0.001 *** |
| Grandprogeny | 0.016 * | 0.001 ** |
| Cumulative sequencing coverage of relatives |  |  |
| Grandparents | 0.456 | 0.297 |
| Parents | 0.245 | 0.021 * |
| Progeny | 0.100 | 0.363 |
| Grandprogeny | 0.044 * | 0.016 * |

<sup>a</sup>LD: low density; HD: high density.
\**p*-value=0.05-0.01; \*\**p*-value=0.01-0.001; \*\*\**p*-value<0.001.

### Factors that affect variant-wise imputation accuracy

The main factors that determined the variant-wise imputation accuracy were the MAF of the variants and the number of sequenced individuals or the cumulative sequencing coverage at the variant site. Whether a marker was present in the marker array or not and the distance of a variant to the nearest variant from the marker array were not influential partitioning factors in the regression trees. The relative position of the variants within the chromosome was used as an influential partitioning factor in the regression tree of the variant-wise genotype concordance but not of the dosage correlation. The results were consistent between the simulated and the real data.

The regression tree for the factors that affect variant-wise dosage correlations on the simulated data is shown in Figure 6a. The first factor that determined variant-wise imputation accuracy was MAF. The imputation accuracy was limited for very rare variants: 0.23 for MAF below 0.001 (n=704), 0.50 for MAF between 0.001 and 0.005 (n=1,217), 0.79 for MAF between 0.005 and 0.028 (n=2,111), and 0.93 for MAF above 0.028 (n=25,968). Other partition factors were the number of individuals with sequencing coverage at a given position, the cumulative sequencing coverage at a given position, and population size. The dosage correlations observed in the real data within each partition of the regression tree followed the same trends as for the simulated data, but ranged from 0.51 (n=11,312) to 0.93 (n=89,701) and were greater than those from the simulated data, especially at low MAF (Figures 6b).

**Figure 6.**
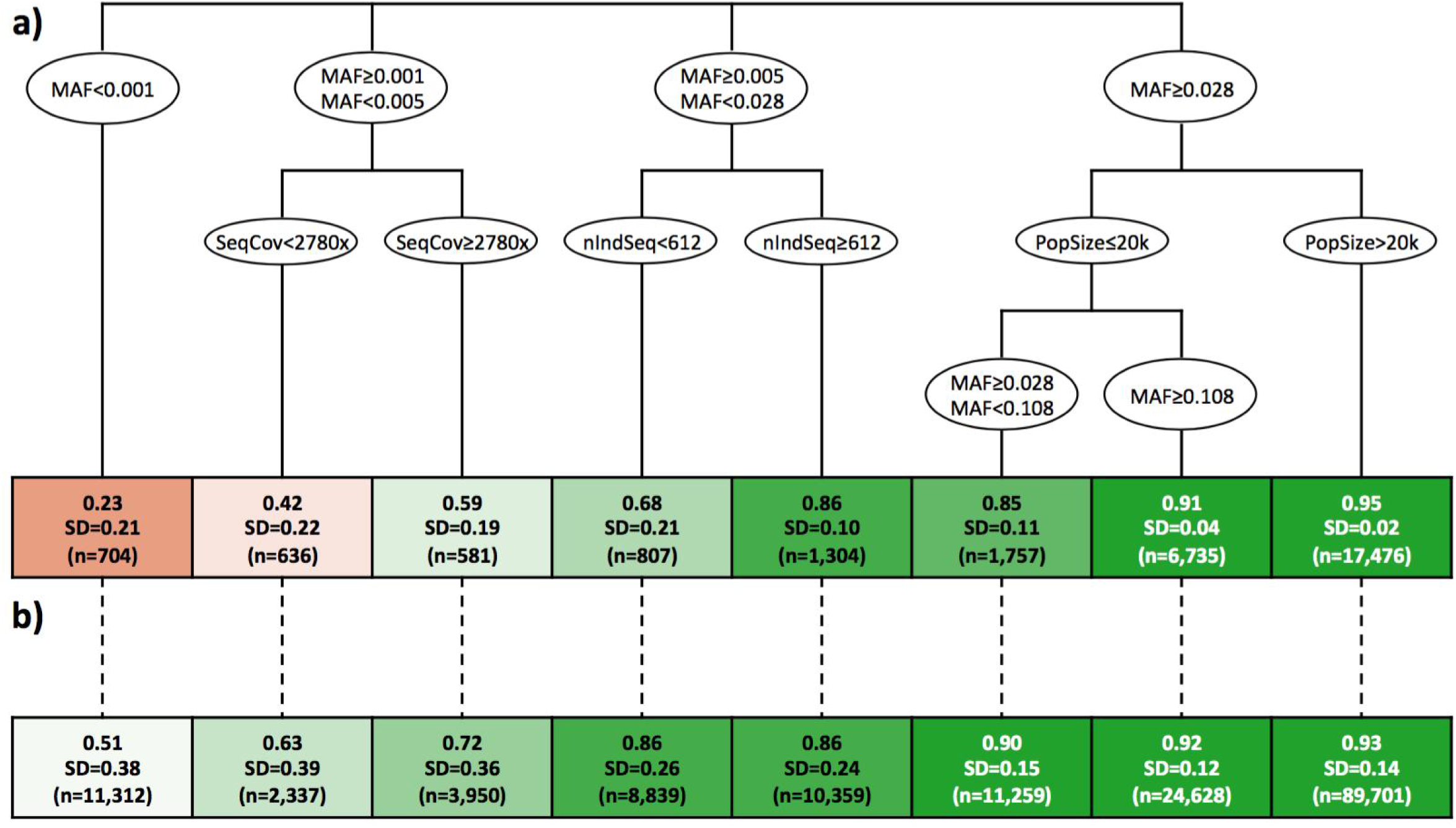
Regression tree of the factors that affected variant-wise dosage correlation in (a) the simulated data and (b) comparison to the real data. Variables include minor allele frequency (MAF), number of individuals sequenced at a position (nIndSeq), cumulative sequencing coverage at a position (SeqCov), and population size (PopSize).

The regression tree for the factors that affect variant-wise genotype concordance in the simulated data is shown in Figure 7a. The genotype concordances showed the opposite trend with MAF than the dosage correlations, with values from 99.0% for MAF below 0.050 (n=5,537) to 93.9% for MAF above 0.154 (n=18,230). For variants with MAF greater than 0.050, the average imputation accuracy increased with the number of individuals that had at least one sequence read covering a given position, from 94.8% (n=1,589) to 97.5% (n=4,644), when MAF was between 0.050 and 0.154, or from 86.1% (n=299) to 95.0% (n=12,668), when MAF was above 0.154. The relative position of the variant within a chromosome was an influential partitioning factor in the case of variants with high MAF and a high number of sequenced individuals. The variants at the extreme ends of the chromosome tended to be imputed with lower accuracy (90.5%; n=152) than those at intermediate positions (94.5%; n=7,786). This variable was not an influential partitioning factor in the regression tree of the dosage correlations. The genotype concordances observed in the real data were consistent with the partitions of the regression tree based on the simulated data (Figures 7b).

**Figure 7.**
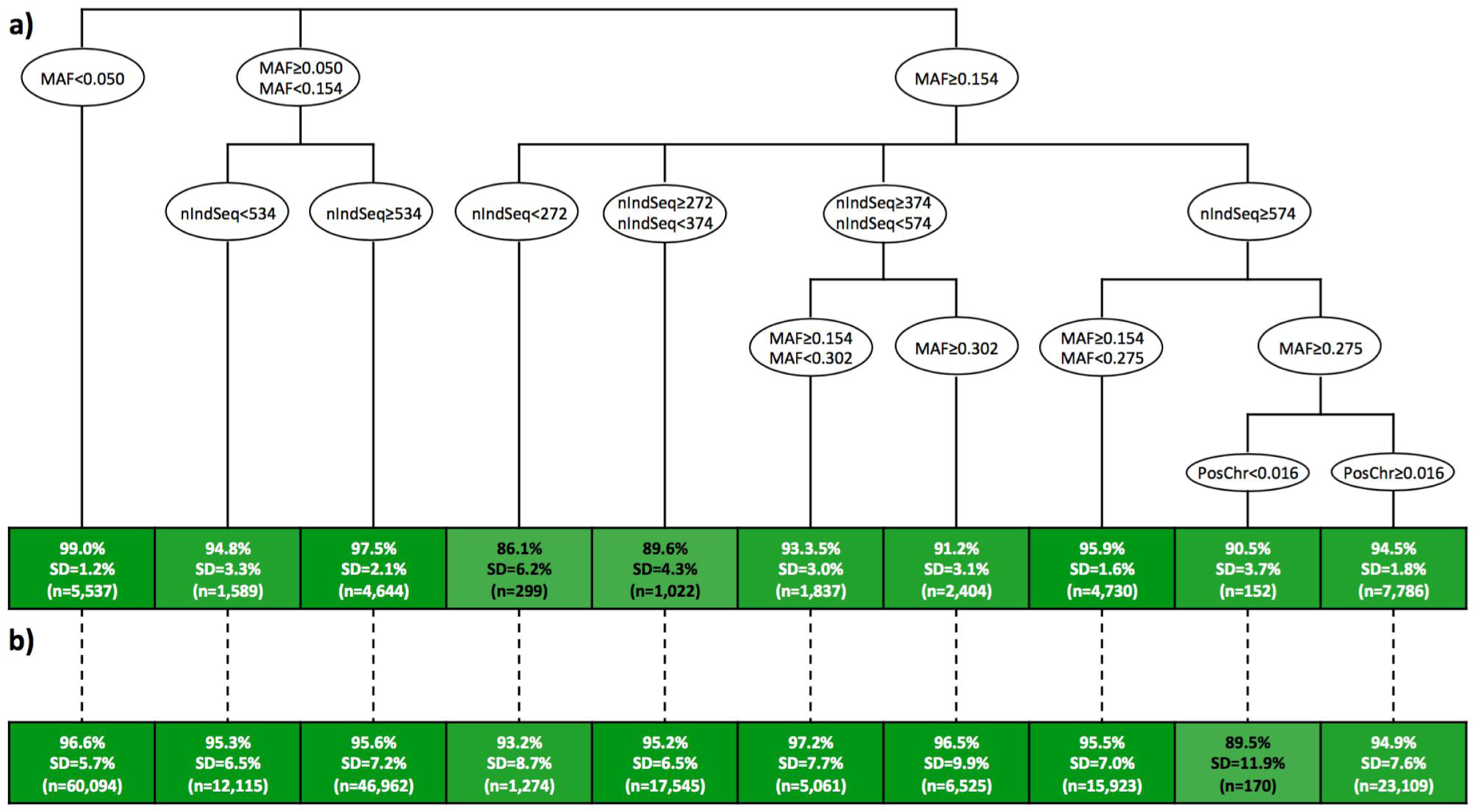
Regression tree of the factors that affected variant-wise genotype concordance in (a) the simulated data and (b) comparison to the real data. Variables include minor allele frequency (MAF), number of individuals sequenced at a position (nIndSeq), and position of the variant within the chromosome (PosChr).

### Impact of data misassignment and pedigree errors

Data misassignment and pedigree errors can have drastic consequences on the imputation results. The impact of data misassignment and pedigree errors, measured as the dosage correlation between the results with and without the deliberate error, is presented in Figure 8 for the target individual (‘ind’) and its immediate relatives. We report here the average dosage correlation but note that there was large case-by-case variability due to the stochasticity of the data misassignment and pedigree errors.

**Figure 8.**
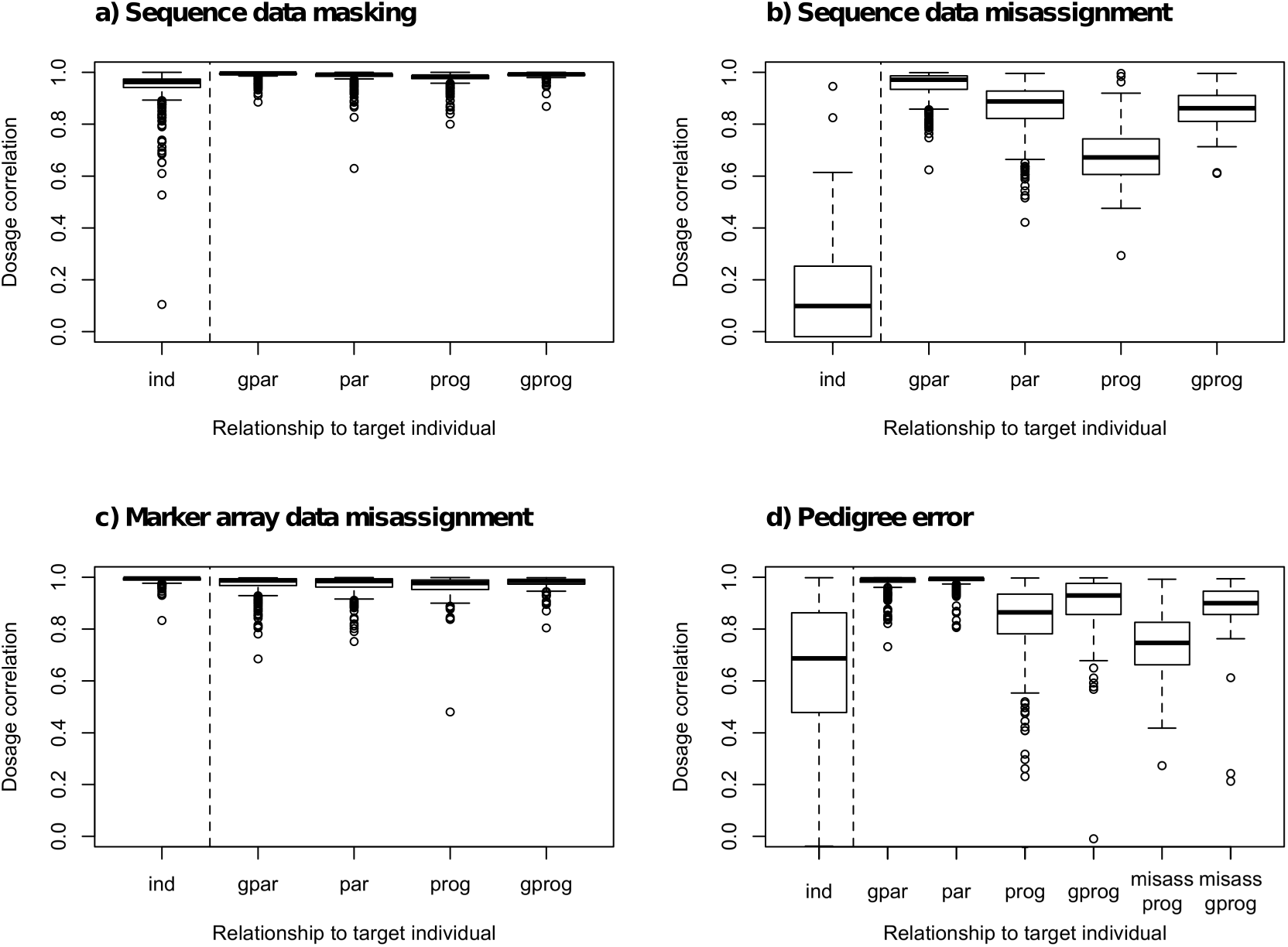
Impact of data misassignment and pedigree errors on imputation accuracy. The dashed line separates the individual directly affected by the data modification (ind) and its relatives (gpar: grandparents, par: parents, prog: progeny, gprog: grandprogeny, misass prog: misassigned progeny, misass gprog: misassigned grandprogeny). The y-axis measures the individual-wise dosage correlation between the imputed genotypes based on complete correct data and either missing or misassigned data for the individual itself and its relatives. In panel (a) we provide the case where the sequence data of the target individual was masked as in Test 1; in panel (b) where the sequence data of another individual was misassigned to the target one; in panel (c) where the marker array data was misassigned; and in panel (d) where we assigned the progeny from one of the individuals sequenced at high coverage to the target individual.

When we removed the high-coverage sequence data of the target individual, as in Test 1 (Figure 8a), the dosage correlation with complete data imputation was 0.94 for the target individual. The impact of removing the sequence data of the target individual had a limited impact on imputing its relatives, which had dosage correlations of 0.97 to 0.99 compared to the case with complete data.

When the sequence data was misassigned (Figure 8b), the dosage correlation of the target individual drastically decreased to 0.13, as did (in order of magnitude) that of its progeny (0.68), then its grandprogeny (0.86) and parents (0.86), and finally its grandparents (0.95).

When the marker array data was misassigned (Figure 8c), the dosage correlation of the target individual remained very high (0.99), probably because the high-coverage sequence data provided high certainty about its true genotypes. Despite this, potential errors in the segregation probabilities resulted in dosage correlations for the relatives of the target individual that were slightly lower (0.97 to 0.98) and showed a greater dispersion.

Finally, when the pedigree was misassigned (Figure 8d), the impact of such errors depended on the number of true and misassigned relatives that the target individual had. In our test the target individual was misassigned progeny from one of the individuals sequenced at high coverage. The dosage correlation of the target individual greatly decreased (0.65). The greatest impact of the pedigree errors was on the misassigned progeny (0.74), but the impact on the true progeny was also large (0.83). The impact was smaller on the misassigned grandprogeny (0.89) and the true grandprogeny (0.90). The dosage correlation of the parents and grandparents of the target individual were largely unchanged (0.99 and 0.98, respectively), probably because they had other correctly assigned relatives (like their own parents) that contributed more accurate data.

## Discussion

In this paper we present the results of a large-scale sequencing study that aimed to generate accurately imputed whole-genome sequence information on hundreds of thousands of individuals. Our results show that we were able to obtain highly accurate sequence information for approximately 230,000 individuals from four different populations that were genotyped at a maximum of 75,000 markers genome-wide, by sequencing only 2% of the individuals in each population, mostly at low coverage. We found that imputation accuracy was high for most individuals, especially for descendants of the first few generations of a pedigree. The same approach was applied to five additional populations (results not shown), providing high-quality whole-genome sequence data for a total of more than 350,000 individuals. To our knowledge this is the largest set of whole-genome sequence information assembled to date in pigs [36] or in any other livestock species (e.g., [7,37]).

Our results give rise to four major points of discussion: (i) the overall performance of the sequencing strategy and the approach that we used for imputing whole-genome sequence data; (ii) the individual-wise imputation accuracy; (iii) the variant-wise imputation accuracy; (iv) the comparison to other imputation methods; and (v) the implications for population-wide sequencing studies.

### Overall performance of the sequencing strategy and hybrid peeling

The overall performance of our sequencing strategy coupled with hybrid peeling was high. We were able to impute whole-genome sequence data for hundreds of thousands of individuals with a median dosage correlation of 0.97 by sequencing only about 2% of the individuals in each of our pedigreed populations. Most of the sequenced individuals were sequenced at low coverage, with 90% of the sequenced individuals at either 1x or 2x and only 6.4% of the sequenced individuals being sequenced at a high coverage of 15x to 30x. Sequencing a subset of individuals at high coverage may improve the variant discovery rates as well as provide a validation set for variants discovered with low-coverage sequence data. It is difficult to separate the contributions of the sequencing strategy and of the imputation method to the imputation accuracy. We have assessed the contribution of the sequencing strategy on imputation accuracy in a companion paper [27]. Overall, sequencing coverage does not seem a very influential factor if a sufficiently large number of individuals is sequenced and, therefore, the sequencing strategy based primarily on low-coverage sequencing that we have described enabled high imputation accuracy in real livestock populations regardless of the size of the population.

Our sequencing strategy and imputation method enabled high imputation accuracies of whole-genome sequence data from marker arrays with relatively low densities, of approximately 15,000 and 75,000 markers genome-wide. The low dependence on marker arrays with higher densities is in contrast to the findings of previous studies on imputation of whole-genome sequence data, which have found that marker array genotyping density was critical when using other sequencing strategies and imputation methods. For example, van Binsbergen et al. [38] found that imputing from marker arrays with a density similar to ours (50,000 markers genome-wide) resulted in low accuracies (dosage correlations of up to 0.80) when using the Beagle imputation software (version 3; [39]) in cattle. Van den Berg et al. [36] found similarly low accuracies in pigs (dosage correlations of around 0.70), probably because the number of sequenced individuals was small. In order to achieve higher imputation accuracies, an intermediate step of imputation to a much higher density (700,000 markers genome-wide or similar) was previously proposed [38]. This intermediate step has been used in several studies and with other imputation methods [36,37,40,41], but this may be a drawback for populations where marker array data at such high densities is not available. We found that a combination of an appropriate sequencing strategy and hybrid peeling achieved high imputation accuracies without any intermediate imputation steps being required for the LD individuals, likely due to the ability of both methods for exploiting pedigree and existing marker array information to maximise the value of the generated whole-genome sequence data for the whole population.

### Individual-wise imputation accuracy

Although most of the individuals had high imputation accuracy, a small portion of individuals had much lower imputation accuracies than the rest. These individuals mostly belonged to the earliest generations of each pedigree. This reduction of imputation accuracy in the earliest generations of the pedigree was consistent with observations in previous simulation studies [15,27]. The individuals involved had very little information available for themselves and for their ancestors, i.e., many of these individuals were not genotyped with marker arrays or their parents and grandparents were not genotyped either. Ancestors are very informative for the phasing of the genotypes and availability of their marker array data determines the accuracy of estimation of the segregation probabilities used in the multi-locus step of hybrid peeling, on which the subsequent single-locus step of hybrid peeling relies.

In a similar way, the marker array density at which the ancestors were genotyped affected imputation accuracy of an individual, regardless of the marker array density at which the individual itself was genotyped. This can be explained by the fact that parental and grandparental genotypes are needed for accurately phasing the individual’s genotype and even a small number of markers suffices to capture the small number of recombinations between the individual and its parents [42]. Thus, strategies that target parents that contribute large number of progeny for genotyping at high density, such as current genotyping practices of breeding programs with genomic selection [43,44], seem appropriate.

Provided that the segregation probabilities were accurately estimated, high connectedness of an individual to the rest of the population enhanced its imputation accuracy by favouring the transmission of information from many relatives and by increasing the likelihood that a closely connected individual has sequence data. In livestock breeding populations, pedigrees are usually deep and individuals have a high degree of relatedness. The connectedness of the imputed individuals to a sufficient number of informative relatives with marker array or sequence data allows for high imputation accuracy (after the initial generations for which the imputation accuracy was low) even when only a small subset of individuals was sequenced at low levels of coverage.

It is critical to perform quality controls of the data before performing imputation to avoid any data misassignment or pedigree errors. In this study we attempted to set an upper threshold for the impact that these errors could have on the individual-wise imputation accuracy of the affected individuals as well as how these errors propagate to the relatives of the affected individuals in a pedigree-based method. We found that the most serious errors occurred due to pedigree errors or assigning sequence data to a wrong individual. However, this may be distorted by the fact that all the target individuals had high-coverage sequence data. Therefore, misassignment of marker array data must not be ignored as it could also have a strong impact on imputation accuracy when it affects individuals that are not sequenced, sequenced at low coverage, or whose relatives are genotyped with low-density marker arrays. Fortunately, frameworks to detect data misassignment [45] and pedigree errors [46] have been developed. We did not test the impact that map errors could have on the imputation accuracy, but it is obvious that they would hamper the estimation of the segregation probabilities and thus imputation accuracy.

### Variant-wise imputation accuracy

We obtained high variant-wise imputation accuracy, especially after filtering out individuals that were likely to have low imputation accuracy. The primary factor for variant-wise imputation accuracy was MAF. This was expected, as MAF is widely known to be one of the main factors that determine imputation accuracy regardless of the imputation method, and we found, similar to other studies, that imputation accuracy was lower for variants with very low MAF [4,38,40,47].

The next most important factors were the total number of reads that covered that variant site and the number of individuals who had sequence data at that variant site. Low-coverage sequencing results in a sparse distribution of reads along the genome, and it is likely that only a subset of the sequenced individuals will have any reads that map to a given variant site and that the cumulative coverage across variant sites will also vary. In our study the number of individuals with some coverage and the cumulative coverage may be confounded because most individuals were sequenced at 1x or 2x, but in general this indicates the importance of having as many sequenced individuals as possible with some coverage at each variant site [27], a circumstance that is favoured by sequencing strategies based on low coverage.

The importance of the number of individuals sequenced at a variant site also suggests that imputation accuracy could be lower in regions with extreme base compositions or particular sequence motifs that hamper read alignment [48,49]. While the complexity of a given region, namely the presence of large repeats, is another factor that could affect local imputation accuracy along a chromosome [40,50], it was not considered in our study.

Inferring the segregation probabilities from the flanking markers that are included in the marker array did not result in noticeably lower imputation accuracy for those variants that were not included in the marker array. Moreover, variant-wise imputation accuracy was found to be independent of the distance between the variant and the flanking markers at which the segregation probabilities were estimated. This is again the reflection of relying on pedigree and the fact that are only few recombinations between a parent and its progeny. However, imputation accuracy tended to be lower for the markers that were at the extreme ends of the chromosome. This affected a relatively small number of variants that were located before the first marker and after the last one and therefore were not flanked on both sides by markers from the arrays. These findings differed from those of previous studies using methods based on linkage disequilibrium (Beagle, version 3; [39]), where variant-wise imputation accuracy decreased as the distance between each variant and the nearest variant in the marker array (from which imputation to whole-genome sequence data was performed) increased [38].

### Comparison to other imputation methods

We did not intend for a direct comparison of the performance of hybrid peeling with other available imputation methods because there are fundamental differences in how they exploit information (pedigree and linkage vs. linkage disequilibrium) and because sequencing strategies and imputation methods are confounded across studies. However, we have previously compared the performance of our hybrid peeling with findhap (version 4; [47]) [15] and other studies have compared other available imputation tools [40,41,47,51], including tools such as Beagle (versions 3 and 4; [39,52]), IMPUTE2 [53], findhap [47], FImpute [54], or Minimac3 [55]. Many of these methods are population-based imputation methods that use an already phased haplotype reference panel to impute genotyped individuals to whole-genome sequence data. As a consequence, previous studies of the factors that influence imputation accuracy have been primarily concerned with the design of the reference panel. Some of these concerns involve the convenience of using single-breed or multi-breed reference panels [41,51], population-specific reference panels [41,56], the availability of marker array data for the sequenced individuals or not (it removes the genotype uncertainty that otherwise would arise from sequencing at low coverage at some pre-established positions) [47], or the trade-off between number of individuals sequenced and sequencing coverage [47]. In contrast, in this paper we used a purely pedigree-based imputation algorithm. This allows us to exploit the large amount of linkage between the haplotypes of an individual and their relatives.

### Implications for population-wide sequencing studies

The coupling of an appropriate sequencing strategy [13,14,27] and an appropriate imputation method, such as hybrid peeling [15], enabled the generation of large datasets of sequenced individuals at a low cost and with high accuracy. This is a critical step for the successful implementation of whole-genome sequence data for genomic predictions, within and across breeds, as well as for fine-mapping of causal variants underlying quantitative traits, which could guide the promotion and removal of alleles by gene editing [57,58].

In this paper we focused on individual-wise imputation accuracy as an indicator of the value of this data for applications such as genomic prediction. Previous studies on imputation accuracy of whole-genome sequence data focused on variant-wise imputation accuracy rather than individual-wise [38,40,47]. In the context of genomic prediction, the estimate of the realized relationship between two individuals will correlate strongly with the individual-wise, but not the variant-wise, imputation accuracy [32,59]. Understanding which factors determine the variability of individual-wise, as well as variant-wise [38,40], imputation accuracy would enable accuracy-aware filtering of the imputed data prior to downstream analyses. With that purpose we used regression trees on simulated data designed to mimic the real data for identifying a small set of partitioning factors that may be used as criteria to filter out individuals with expected low imputation accuracy.

## Conclusion

We used hybrid peeling to impute whole-genome sequence data of hundreds of thousands of individuals from real livestock populations that were genotyped at a maximum of 75,000 markers genome-wide by sequencing only 2% of the individuals of each population, mostly at low coverage. The coupling of an appropriate sequencing strategy and hybrid peeling is a powerful method for generating whole-genome sequence data in large pedigreed populations, as long as the individuals are connected to enough informative relatives with marker array or sequence data, and regardless of population size. The characterization of the factors that affect the individual-wise and variant-wise imputation accuracy of hybrid peeling can inform genotyping and sequencing strategies as well as provide accuracy-aware quality control guidelines for the imputed data before downstream analyses. The success of this sequencing strategy demonstrates the possibility of obtaining low-cost whole-genome sequence data on large pedigreed livestock populations, which is a critical step for the successful implementation of whole-genome sequence data for genomic predictions and fine-mapping of causal variants.

## Ethics approval and consent to participate

The samples used in this study were derived from the routine breeding activities of PIC.

## Consent for publication

Not applicable.

## Availability of data and material

The datasets generated and analysed in this study are derived from the PIC breeding programme and not publicly available.

## Authors’ contributions

RRF, AW, and JMH designed the study; RRF and CYC performed the analyses; RRF wrote the first draft; AW, GG, WOH, AJM, and JMH assisted in the interpretation of the results and provided comments on the manuscript. All authors read and approved the final manuscript.

## Acknowledgements

This work has made use of the resources provided by the Edinburgh Compute and Data Facility (ECDF) (http://www.ecdf.ed.ac.uk/).

## Supplementary Information

**Figure S1.**
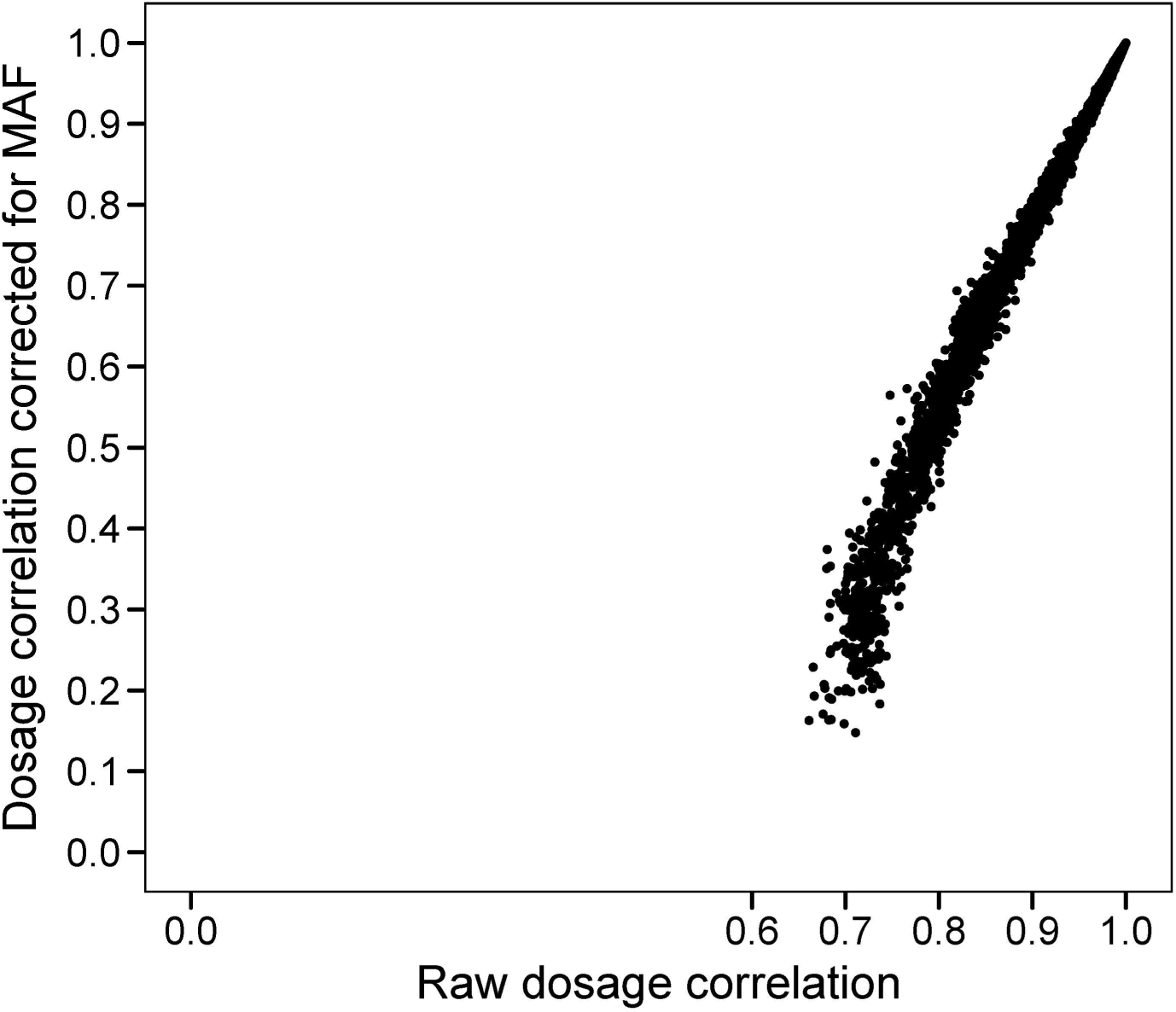
Relationship between raw and MAF-corrected individual-wise dosage correlations for sequence data. Results are for simulated data with a pedigree with 30k individuals an investment equivalent to 2% of the population sequenced at 2x.

**Figure S2.**
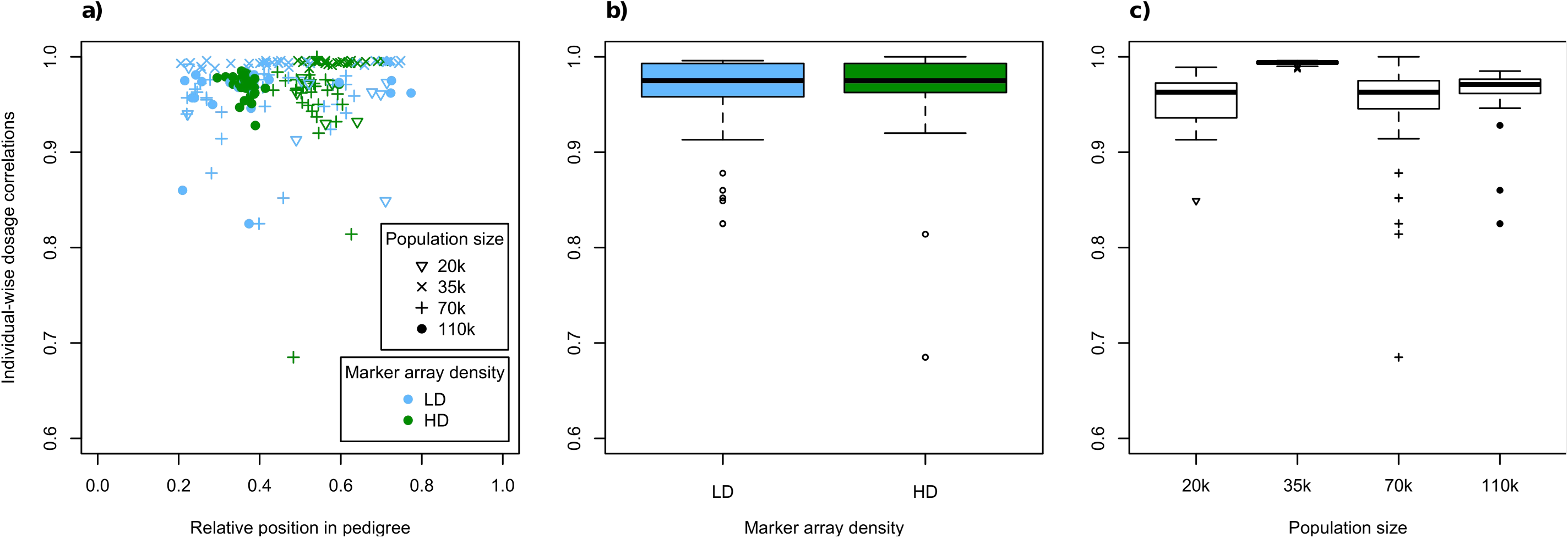
Individual-wise dosage correlation on the real data after excluding the individuals in the first 20% of the pedigree with respect to (a) relative position of the tested individuals within a pedigree, (b) genotyping marker array density, and (c) population size.

**Figure S3.**
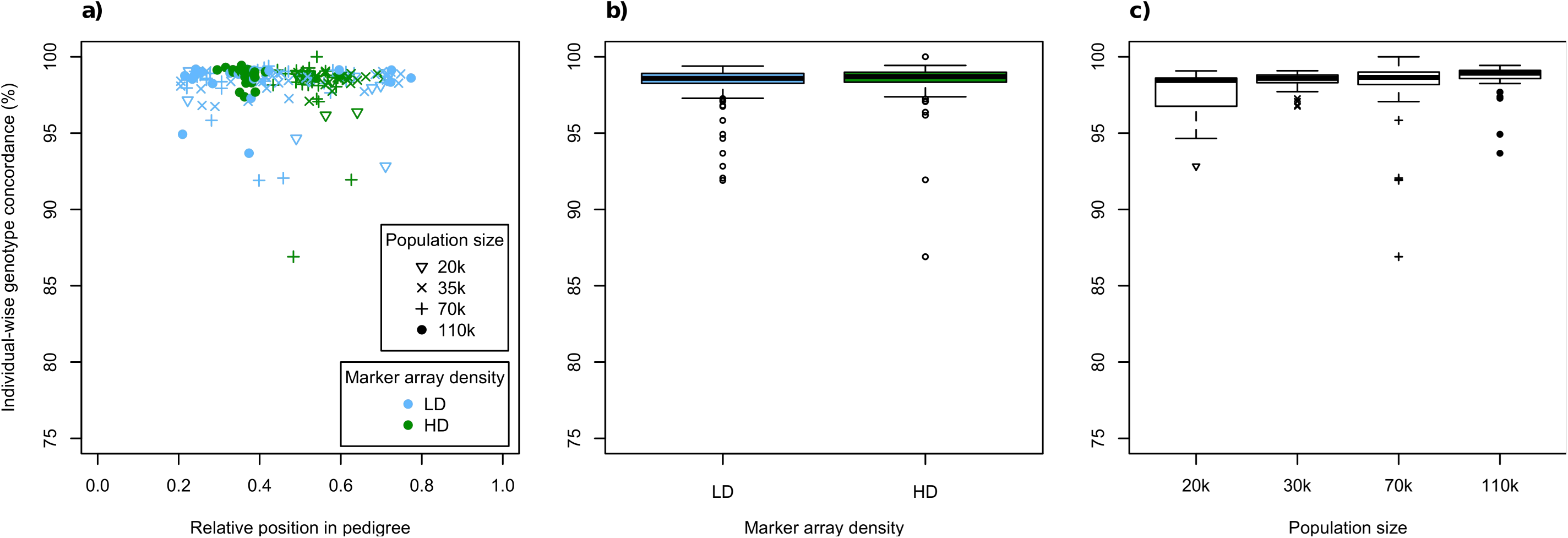
Individual-wise genotype concordance on the real data after excluding the individuals in the first 20% of the pedigree with respect to (a) relative position of the tested individuals within a pedigree, (b) genotyping marker array density, and (c) population size.

**Figure S4.**
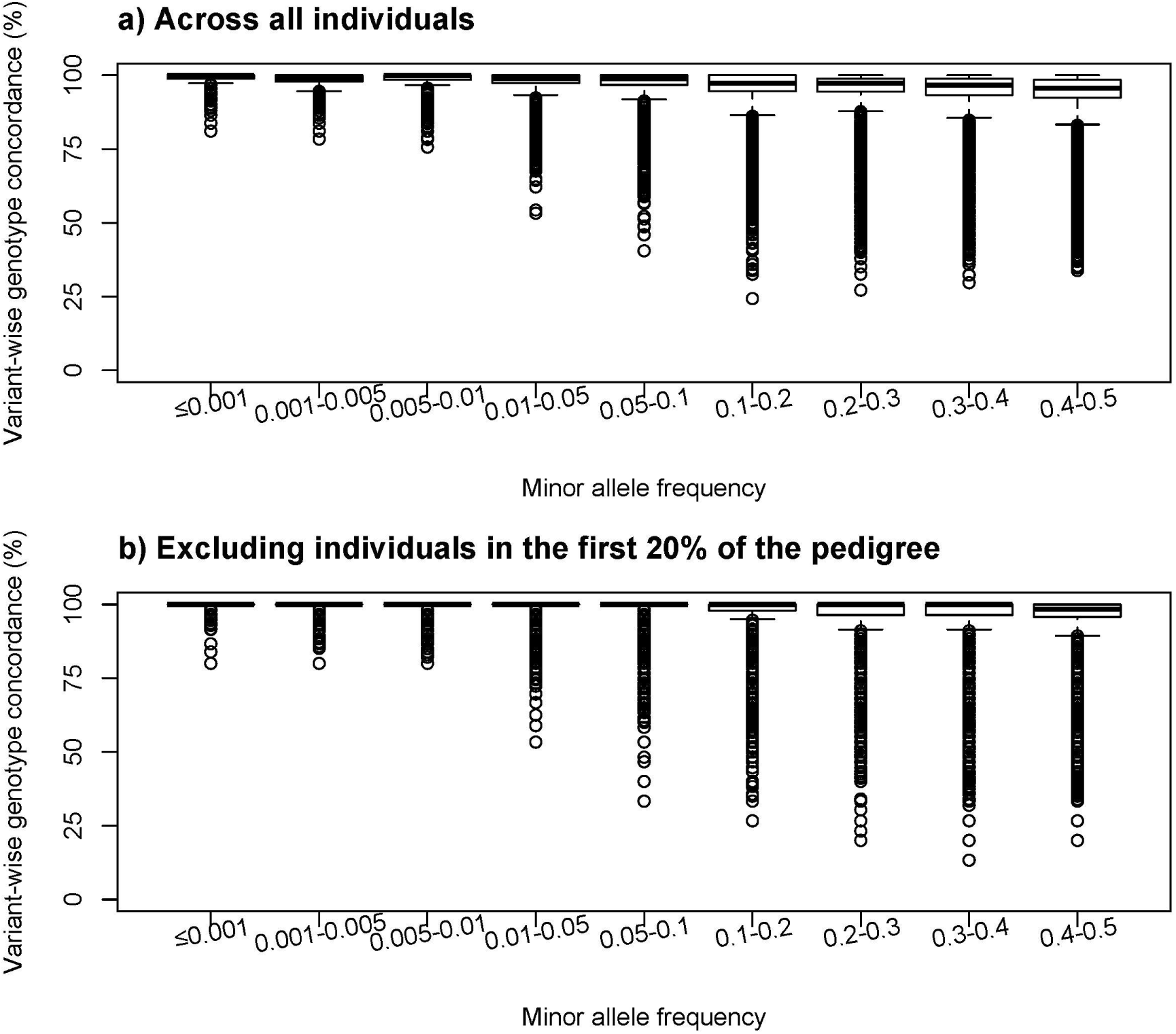
Variant-wise genotype concordance on the real data respect to minor allele frequency. Results are shown for (a) all individuals or (b) after excluding the individuals in the first 20% of the pedigree.

